# G4-Ligand Conjugated Oligonucleotides Mediate Selective Binding and Stabilization of Individual G4 DNA Structures

**DOI:** 10.1101/2023.09.20.558437

**Authors:** Andreas Berner, Rabindra Nath Das, Naresh Bhuma, Justyna Golebiewska, Alva Abrahamsson, Måns Andréasson, Namrata Chaudhari, Mara Doimo, Partha Pratim Bose, Karam Chand, Roger Strömberg, Sjoerd Wanrooij, Erik Chorell

**Author notes:** Equally contributing corresponding authors **Corresponding Authors** Sjoerd Wanrooij, Erik Chorell. Equally contributing first authors (Andreas Berner, Rabindra Nath Das, Naresh Bhuma).

## Abstract

G-quadruplex (G4) DNA structures are prevalent secondary DNA structures implicated in fundamental cellular functions such as replication and transcription. Furthermore, G4 structures are directly correlated to human diseases such as cancer and have been highlighted as promising therapeutic targets for their ability to regulate disease-causing genes, e.g., oncogenes. Small molecules that bind and stabilize these structures are thus valuable from a therapeutic perspective and helpful in studying the biological functions of G4 structures. However, there are hundreds of thousands of G4 DNA motifs in the human genome, and a longstanding problem in the field is how to achieve specificity amongst these different G4 structures. Here, we have developed a strategy to selectively target an individual G4 DNA structure. The strategy is based on a ligand that binds and stabilizes G4s without selectivity, conjugated to a guide oligonucleotide, that specifically directs the G4 Ligand conjugated Oligo (GL-O) to the single target G4 structure. By employing various biophysical and biochemical techniques, we show that the developed method enables the targeting of a unique, specific G4 structure without impacting other off-target G4 formations. Considering the vast amounts of G4s in the human genome, this represents a promising strategy to study the presence and functions of individual G4s but may also hold potential as a future therapeutic modality.

## INTRODUCTION

G-quadruplex structures (G4s) are non-canonical four-stranded DNA structures observed in guanine-rich regions of genomes. These structures are composed of square planar arrangements of four guanines (G) that are held together by eight Hoogsteen hydrogen bonds to form G-tetrads. A G4 structure is typically composed of two to four G-tetrads stacked on top of each other and can be highly stable and polymorphic (1, 2). The direction of the G-rich strands and the connecting loops can give the G4 structures different conformations (2). However, the core of G4 structures with stacked G-tetrads is always the same.

Computational studies have revealed around 700,000 sequences in the human genome with potential to form G4 structures (3-6). These sequences showed a non-random distribution with G4 motifs located predominantly at e.g., the promoter regions of oncogenes (4, 7). G4 DNA structures are mainly formed in single-stranded G-rich sequences in the course of unwinding of the double helix during different biological processes such as; replication, transcription, translation, DNA repair, molecular crowding, and negative supercoiling (8-10). G4 structures are involved in regulating many cellular processes such as telomeric length maintenance and transcription but can also be linked to human diseases such as neurodegenerative diseases,(11, 12) and different types of cancers(13, 14). For example, the *c-MYC* gene, that is involved in cell cycle regulation and is overexpressed in most cancer types (15, 16), is predominantly regulated by the guanine-rich promoter region (NHEIII) Pu27 which can fold into multiple G4 structures (17, 18).

The ability to silence oncogenes by stabilizing G4 structures in their promotors, like the *c-MYC* G4, has appeared as an attractive novel therapeutic approach and especially for those oncogenes coding for “undruggable” proteins (13). Indeed, there are vast examples of small organic molecules developed to bind and stabilize G4 DNA with selectivity over double-stranded DNA (14, 19-21). In the cellular context, such ligands have also been observed to down-regulate the transcription of oncogenes (22). In the same way, telomerase activity has been found to be inhibited by ligands that can stabilize the telomeric G-quadruplex structures (23, 24). These types of compounds can thus be used to study G4 biology and for further developments towards therapeutics. However, the quantity of G-quadruplex ligands that proceed into clinical trials is remarkably low. Compound CX3543 is one example that entered clinical studies but was eventually withdrawn and its development was discontinued (13, 25). An analogue based on this compound, CX-5461, is now in phase I/II clinical trials for patients with BRCA1/2 deficient tumors (26, 27). The major reason for the low progression towards clinical trials is likely related to the lack specificity, as most G4 ligands can bind to several G4 structures leading to a plethora of side effects considering the hundreds of thousands of G4 motifs in the human genome.

The need for G4 DNA stabilizing compounds able to uniquely recognize only one G4 structure has been highlighted for decades but still with limited to no compounds with confirmed specificity. This is mainly ascribed to the fact that the core of G4s is the same between different structures and offers a flat and hydrophobic surface that is easily targeted with small molecules. However, this binding mode makes it problematic to discriminate between different G4 structures. In this regard, different approaches to gain selectivity have recently started to be explored such as ligand-peptide conjugates (28), simultaneous recognition of duplex and quadruplex motifs (29, 30), a DNA molecule that hybridize with the flanking single strand to target RNA G4s (31), and ligand-PNA conjugates (32).

To meet the need for G4 specific targeting approaches, we here use the fact that most G4-ligands bind the terminal G-tetrad of the G4 DNA structure. This enables conjugation of the G4-ligand to an oligonucleotide that base-pair with the sequence directly flanking the G4 structure. These G4 flanking regions are heterogenous and differ between each G4, thus allowing specific targeting of individual G4s (Figure 1A). Hence, we have conjugated two recognition motifs; a G4-ligand that targets the terminal G-tetrad of the G4 and a DNA oligonucleotide complementary to the sequence flanking the target G4. The fundamental biochemical process of hybridization between complementary DNA strands into a duplex will guide the G4 ligand-conjugated oligo (GL-O) moiety to the targeted G4 structure. Our hypothesis being that a robust effect, minimizing off-target interactions, is achieved only when both these two recognition motifs can bind simultaneously. We examined this strategy on the Pu24T G4 structure which is a well-known model system of the *c-MYC* oncogene Pu27 (33). To challenge the specificity gained by the GL-O approach, different biophysical and biochemical techniques were employed. Taken together, the results show that the developed approach allows for targeting of a distinct, individual G4 structure while leaving other off-target G4s unaffected. We thus developed an approach with potential to modulate gene expression at the DNA level, which in theory could be designed towards any of the hundreds of thousands of potential G4 DNA structures by replacement of the guide oligonucleotide. Considering the significant developments and successful clinical trials leading to approvals of dozens of oligonucleotide-b ased therapeutics over the last years, the proposed strategy may hold promise for future therapeutic approaches but also has potential as a valuable tool to explore the cellular function of one specific G4 structure.

**Figure 1.**
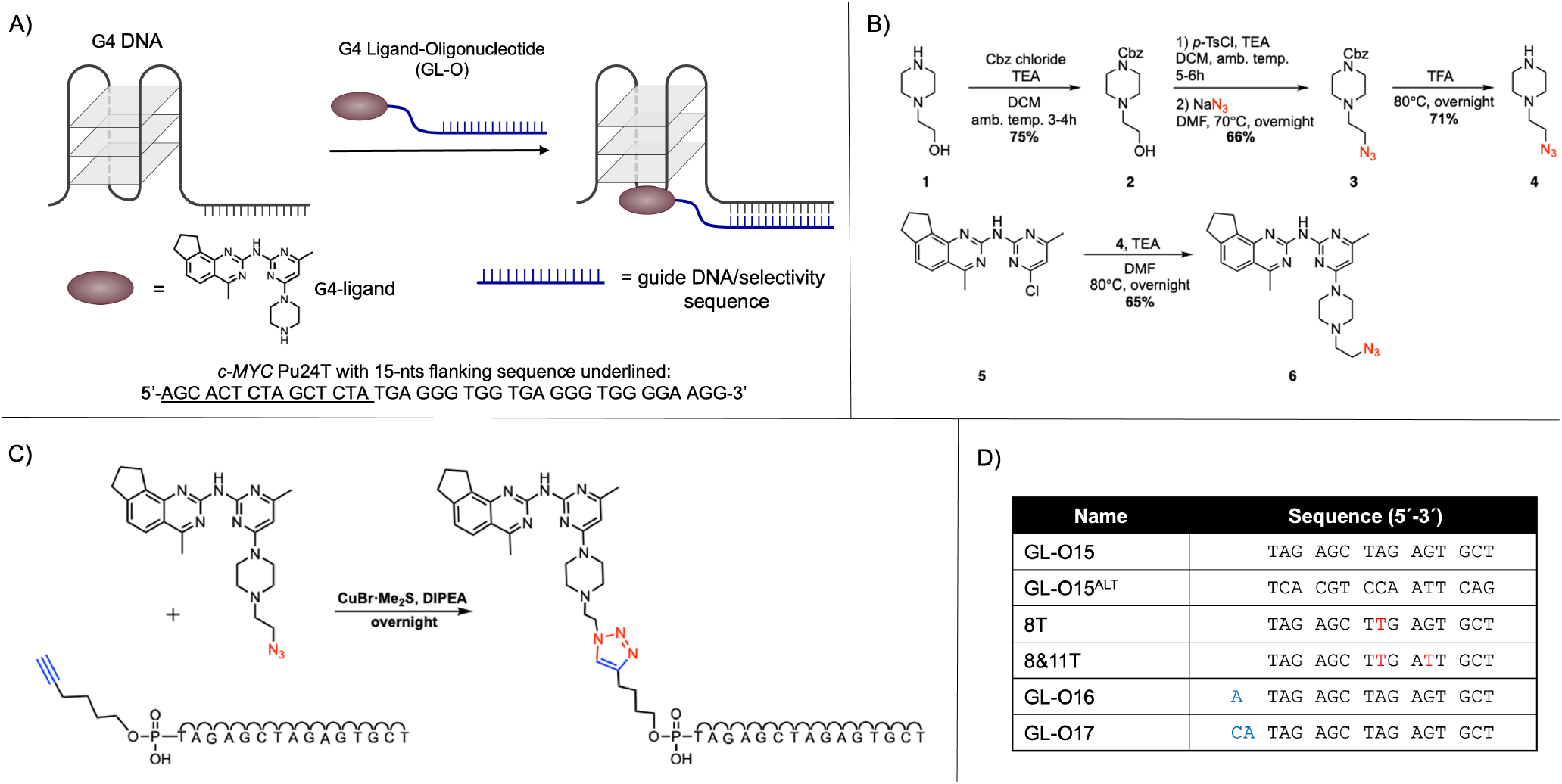
Overview of the strategy and chemistry employed in this study. **A)** Summary of the presented strategy to target individual G4 DNA structures using G4 ligand-oligonucleotide (GL-O) compounds. **B)** Synthesis of the azido functionalized quinazoline-pyrimidine ligand for further conjugations with click chemistry. **C)** Conjugation through copper catalysed click chemistry of the G4-ligand (GL) to the oligonucleotide to give the GL-O15. **D)** Overview of some GL-Os used in this study, all compounds are shown in Table S1.

## RESULTS AND DISCUSSION

### Design and synthesis of G4-ligand conjugated oligonucleotides (GL-Os)

We recently reported on a novel G4-ligand (compound no 14a(34)), that both efficiently binds and stabilizes G4 DNA structures and that has drug-like properties (e.g. no permanent charges and has a low molecular weight) allowing further conjugation to an oligonucleotide guide sequence. To enable conjugation of this G4-Ligand (in this study renamed GL) to the oligonucleotide sequence, an azido substituted piperazine side chain was introduced for further conjugation through click chemistry (Figure 1B). The azido-piperazine (**4**) was synthesized starting from the commercially available 2-hydroxyethyl piperazine **1**, which was protected as its benzoylformate (Cbz) derivative **2** using benzoyl chloroformate (CbzCl). The primary hydroxyl group was next converted into its tosyl derivative which was subsequently displaced with an azido group using sodium azide to get the piperazine-ethylazido derivative **3** in 66% yield. The Cbz group in **3** was deprotected using TFA at 80 °C, to give the desired piperazine-ethylazido linker **4** in 71% yield. A substitution reaction with this linker (**4**) and the chloro-substituted quinazoline-pyrimidine G4-ligand **5** gave the desired azido-functionalized G4-ligand **6** in 65% yield.

After the synthesis of the azido-functionalised G4-ligand, we performed the conjugation step with terminal alkyne modified oligodeoxynucleotide sequences using a copper-mediated click reaction (Figure 1C and Table S2). This enables the production of GL-Os with different guide sequences. (Figure 1D and Table S1).

To test the G4-Ligand conjugated Oligo (GL-O) strategy, we targeted the *c-MYC* Pu24T G4 which is a parallel G4 structure frequently used as model to study the *c-MYC* G4 structure (33). In our experiments with *c-MYC* Pu24T, we also included a 15nts oligonucleotide G4-flanking sequence with AT/GC ratio close to one (1.14)(Figure 1A). This particular 15-nts was selected as the G4-flanking region, because circular dichroism (CD) spectroscopy showed that it did not alter the parallel G4 structure topology of *c-MYC* Pu24T. Other G4-flanking sequences were excluded based on CD evaluation since they altered the topology of the *c-MYC* Pu24T G4 structure (Figure S1A).

## The G4-Ligand Maintains G4 Binding Capacity After Guide Oligonucleotide Conjugation

There are today many reported G4-ligands with high affinity and the main aim of this study is not to improve these but rather to retain the affinity and gain specificity. Therefore, we first investigated if the conjugation of the G4-ligand to a comparatively large oligonucleotide would affect the G4 binding ability by comparing the G4-ligand (GL) alone with the synthesized G4-Ligand conjugated Oligonucleotide (GL-O) in a series of biophysical and *in vitro* biochemical methods.

With nuclear magnetic resonance (NMR) we can simultaneously study the G4 (*c-MYC* Pu24T) and duplex DNA formation (annealing of guide oligonucleotide to the 15-nts G4-flanking region). G4 imino signals are found between 9-12 PPM while the double stranded DNA signals appear between 12-14 PPM. Thus, the co-existence of double-stranded DNA and G4 DNA can easily be monitored in the same NMR spectrum. We recorded ^1^H-NMR spectra of pre-folded *c-MYC* Pu24T G4 alone and in the presence of GL, GL-O15, and O15 (the guide oligonucleotide without G4-ligand) (Figure 2A and S2B). The ^1^HNMR spectrum of *c-MYC* Pu24T G4 structure with the 15-nts G4-flanking sequence was first recorded and the G4 imino protons (10.5-11.8 ppm) display a similar pattern to Pu24T without the inserted G4-flanking sequence, which again confirmed that the added 15-nts does not substantially affect the fold of the G4 structure (Figure S2A). When one equivalent of O15 is added, new dsDNA signals are observed in the 12.3-13.8 ppm region, indicating hybridization of O15. (Figure 2A). The G4 imino signals are unchanged upon O15 addition, showing that the pairing of O15 at the flanking region does not substantially affect the structure or conformation of the *c-MYC* Pu24T G4 structure. Addition of either GL or GL-O15 changes the G4-imino signals (9-12 PPM) showing ligand-G4 structure interaction but the dsDNA signals (12-14 PPM) were only observed with GL-O15 (Figure 2A and Figure S2B). The 15-nts flanking sequence thus seem to serve as recognition sequence able to direct the GL-O15 without disturbing the ability of the GL to bind the G4 structure. CD spectroscopy of *c-MYC* Pu24T confirmed our conclusions from NMR analysis; after addition of GL-O15 we still see the characteristic parallel G4 peaks but also a shoulder at 283-290 nm fitting with dsDNA formation (Figure S3A).

**Figure 2.**
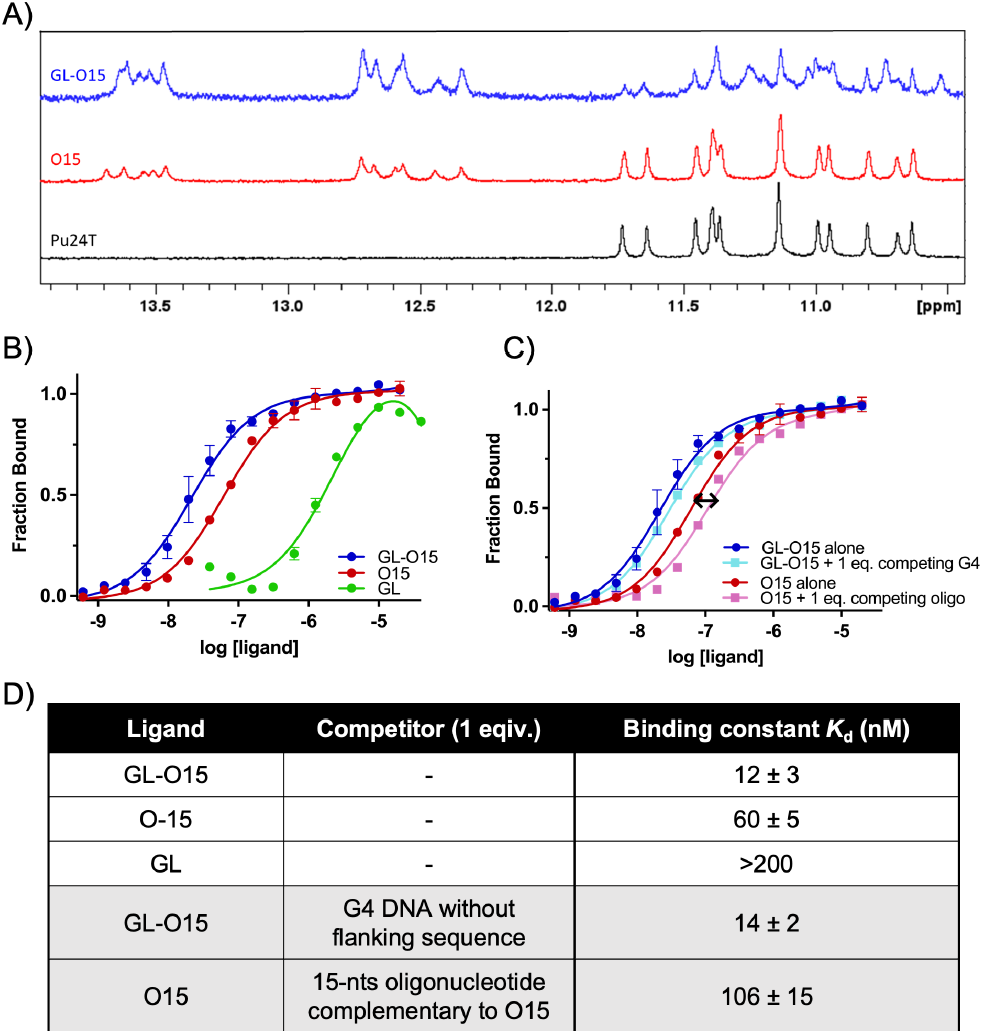
Conjugation of the G4-ligand to the Guide oligonucleotide retain G4 binding. The recognition sequence guide the ligand without affecting G4 binding. **A)** ^1^H NMR spectra of *c-MYC* Pu24T G4 DNA (90 μM) with the flanking sequence (Black), in presence of 1 equivalent of O15 (red) and GL-O15 (blue). G4 imino signals appear between 9-12 PPM and double stranded DNA signals between 12-14 PPM. **B)** Dose-response curves obtained from MST analysis on *c-MYC* Pu24T G4 DNA with the flanking sequence after addition of GL-O15, O15, and GL. **C)** Dose response curves obtained from MST analysis on *c-MYC* Pu24T G4 DNA with the flanking sequence after addition of GL-O15 + 1 equivalent *c-MYC* Pu24T G4 DNA without the flanking sequences as competitor), O15 + 1 equivalent 15-nts oligonucleotide complementary sequence to O15 as competitor. **D)** *K*_*d*_ values obtained from MST analysis.

We next used Microscale Thermophoresis (MST) to determine the binding affinity of GL, O15, and GL-O15 to the *c-MYC* Pu24T G4 structure including flanking sequence (Table S1) (Figure 2B-D). The GL has a binding affinity constant, *Kd*, above 200 nM whereas the O15 display an affinity of 60 nM (Figure 2B and 2D). Encouragingly, the GL-O15 compound has a *Kd* of 12 nM which is considerably stronger than both the GL and the O15 alone. This show that the conjugation of the GL to the oligonucleotide does not negatively affect the binding affinity but on the contrary seem to result in a synergistic binding effect (Figure 2B and 2D). Furthermore, GL-O15 retain its binding affinity to the G4 with the flanking sequence, even in the presence of one equivalent of competing unlabelled *c-MYC* Pu24T G4 DNA without flanking region (Figure 2C and 2D). Binding of the O15 oligo alone, however, was easily out competed by addition of one equivalent of an oligonucleotide complementary to O15 (Figure 2C and 2D).

### Conjugation of the G4-ligand to the Guide Oligonucleotide Retains G4 Stabilization

With the knowledge that GL conjugation to the oligonucleotide did not negatively affect G4 binding but instead resulted in a synergistic improvement, we next investigated if GL conjugation to the oligonucleotide would also retain its ability to stabilize G4s using DNA melting experiments. The CD melting studies with *c-MYC* Pu24T in the presence of O15 showed a stepwise denaturation with increasing temperatures, where the dsDNA (O15 hybridized to the 15-nts flanking region) is denaturing first followed by the G4 DNA structure (Figure 3A-B, S3B-C). Addition of GL-O15 results in an increased melting temperature, which show that the strong binding of GL-O15 to *c-MYC* Pu24T also translates into stabilization of the G4 DNA structure (Figure 3C). In this assay, the increase in melting temperature was slightly lower compared (Figure S1B) to the experiment with addition of GL alone (without O15). The unbound oligonucleotide of GL-O15 thus prevents the conjugated G4-ligand to stabilize the G4 to its full potential. However, this effect is likely linked to the increased temperature in this assay and the fact that the dsDNA is denaturing at a lower temperature compared to the G4 structure. This could be confirmed by NMR experiments at increased temperatures, which show that the dsDNA peaks (12-14 PPM) disappear around 50°C and the G4 DNA signals (9-12 PPM) around 55°C (Figure 3D). Interestingly, the dsDNA peaks are still visible at higher temperatures in the presence of GL-O15 (55°C) compared to when O15 alone is added (50°C), showing that GL-O15 binding to the G4 DNA is also able to stabilize the oligonucleotide hybridization to the 15-nts flanking region.

**Figure 3.**
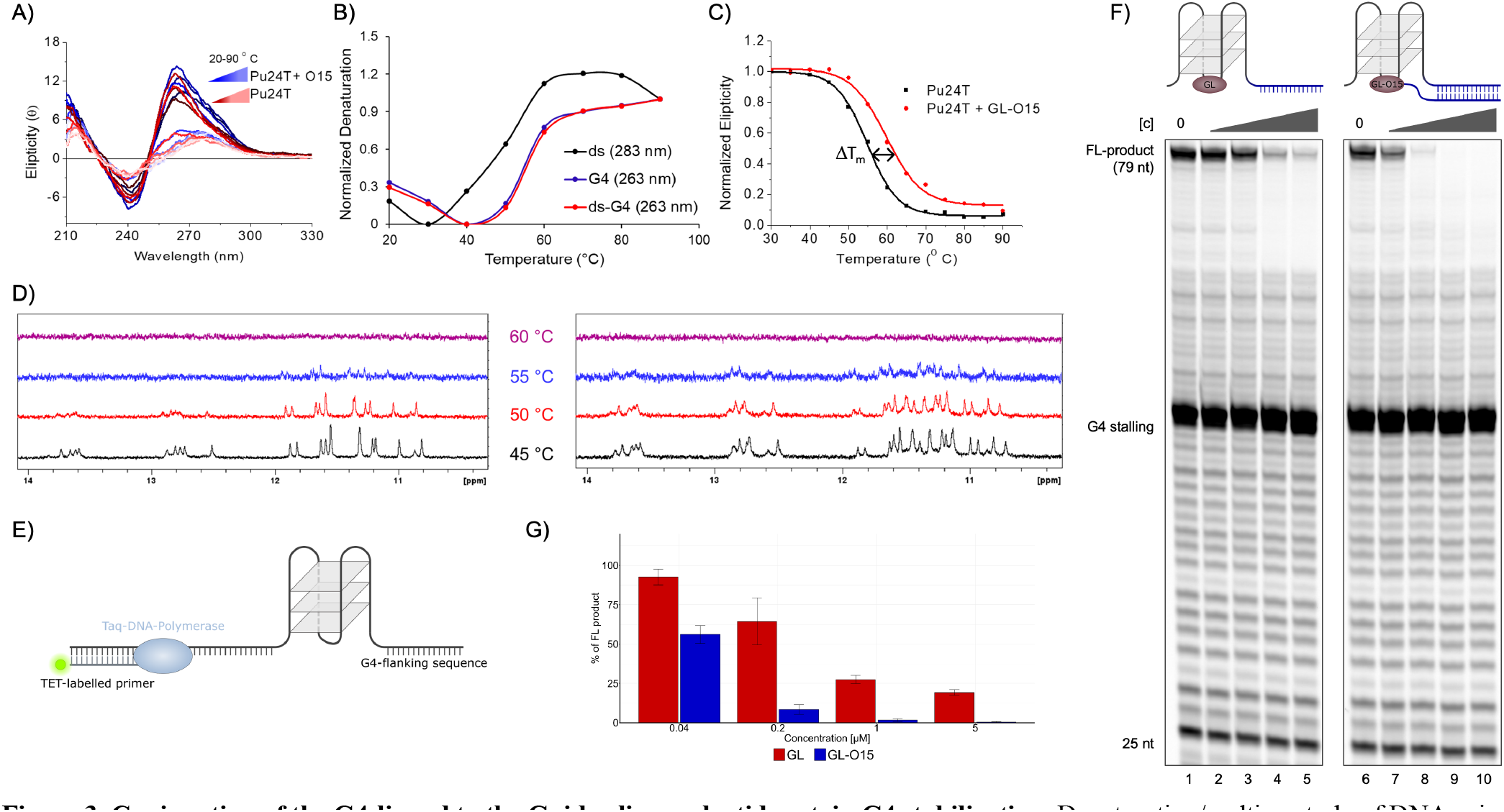
Conjugation of the G4-ligand to the Guide oligonucleotide retain G4 stabilization. Denaturation/melting study of DNA using CD and NMR. **A)** CD spectra of Pu24T in absence of (red gradient) and in presence of O15 (blue gradient) at different temperatures. **B)** Comparison of melting temperature of double stranded (ds), G4, and ds-G4 DNA. **C)** CD melting curve of Pu24T in presence and absence of GL-O15. **D)** ^1^H-NMR spectra of *c-MYC* Pu24T at different temperatures with O15 (left) or GL-O15 (right). **E)** Schematic representation of the Polymerase-Stop-Assay. The TET-labelled primer (grey) is annealed to a G4 forming DNA template (blue) and extended by TaqPolymerase (light blue). **F)** Taq-Polymerase Stop assays in the presence of increasing concentrations (0.04, 0.2, 1, and 5 μM) of GL (lanes 2-5) or GL-O15 (lanes 7-10). **G)** Quantification of the Taq-polymerase Stop assays in F. The full-length product is expressed in % of the full-length band intensity that was obtained in the control reaction that did not contain compound. Mean and standard deviation of three individual experiments is shown.

To further explore how the high affinity of the GL-O15 translates into G4 stabilization, we next used a DNA polymerase stop assay. This assay evaluates how efficiently the Taq polymerase can synthesize new DNA by copying a template strand and give a single nucleotide resolution readout by extending a fluorescent 5’ end-labelled DNA primer. A G4 structure is a hurdle for the DNA polymerase, and if a G4 DNA structure is used on the template strand, the Taq polymerase will stall some nucleotide(s) before the G4 structure. However, the DNA polymerase can partly bypass the G4 structure and synthesize to the end of the DNA template producing a full-length (template run-off) DNA product. The proficiency of G4 stabilizing compounds can be determined by their capability to increase DNA polymerase pausing at the G4 structure, consequently reducing the amount of full-length DNA product.

The experiments were carried out on synthetic G4 forming DNA templates that includes a 15-nts G4-flanking DNA sequence that was complementary to the guide oligonucleotide sequence of GL-O15 (Figure 3E-G). The DNA substrates were designed such that the DNA polymerase encounters the G4 structure at the 19^th^ base after initiation of DNA synthesis from the 25-nts long TET-labelled primer (Figure 3E).

The DNA polymerase stop assays were performed in the presence of increasing amount of either GL or GL-O15 and the G4 stabilizing effect was measured by quantification of the fulllength product (Figure 3F-G). When GL-O15 is presented with a substrate that contains the complementary sequence it results in a strong and dose-dependent G4 stabilization, resulting in a strongly reduced formation of full-length product (Figure 3F lanes 6-10). Compared to the thermal CD melting experiments, this more biologically relevant assay also shows that GL-O15 display an increased G4 stabilizing ability compared to GL (Figure 3G and 3F compare lane 2-5 with 7-10). This shows that base-pairing of the guide oligonucleotide sequence of GL-O15 to the DNA template in the vicinity of the G4, which primarily serves to guide the ligand to the target, simultaneously enhances its G4 stabilizing capacity.

We next performed DNA polymerase stop assays to investigate if GL-O15 induced reduction of full-length product (as shown in Figure 3G and 3F lanes 7-10 and Figure S4 lanes 16-20) is the additive effect of stalling induced by the oligonucleotide annealing and G4-ligand stabilization. Addition of oligo alone (O15) did reduce the full-length product, a direct consequence of the DNA polymerase running into the annealed oligonucleotide which generates an oligonucleotide stalling site (Figure S4 lanes 2-5). The reduction of full-length products is substantially stronger when both O15 and GL are added separately to the same reaction (Figure S4, compare lanes 12-15 with lanes 2-10), most likely due to the additive inhibition of DNA polymerization by 1) the annealed oligonucleotide induced stalling and 2) the G4 stabilization. Interestingly, addition of GL-O15, where the G4-ligand is chemically linked to the guide oligonucleotide results in an even stronger reduction of full-length DNA product suggesting a synergistic effect on G4 stabilization (Figure S4, lanes 17-20). In conclusion, the DNA polymerase stop experiments agree with the biophysical analysis and show that GL conjugation to the oligonucleotide (to form the GLO15) does not only retain the stabilization effect from the GL but results in a synergistic effect that enhances stabilization of the correct G4 structure and prevent stabilization of off-target G4 structures.

To evaluate whether our approach is applicable to G4s beyond the *c-MYC* G4, we also examined the *c-KIT* and *HelB* G4 structures in primer extension assays (Figure S5). Analogous to our findings with the *c-MYC* G4, GL-O15 effectively enhanced G4 stabilization for the *c-KIT* and *HelB* G4s when a complementary flanking region was present on the DNA template. A much stronger reduction of full-length DNA product was observed in the presence of GL-O15 compared to GL for both G4s (*c-KIT*, compare Fig S5 lanes 2-5 with 7-10; *HelB* compare Fig S5 lanes 17-20 with 22-25). This shows that our GL-O strategy in theory can be designed towards any G4 DNA structure by replacement of the guide oligonucleotide.

### The Guide Oligonucleotide Induce G4-ligand Selectively for an Individual G4 Structure

To prove that the increased ability of GL-O15 to bind and stabilize the target G4 is specific, we first performed experiments with c-MYC Pu24T that does not contain a complementary flanking region and thus is unable to hybridize with the oligonucleotide (O15) sequence of GL-O15. Encouragingly, we were unable to detect any binding of GL-O15 to c-MYC Pu24T G4 in the absence of a complementary flanking sequence and a strongly reduced binding using a non-complementary flanking sequence (Figure 4A). To confirm that this selectivity also translates into G4 stabilization, we used the DNA polymerase stop assay with a DNA template containing an alternative (noncomplementary) flanking region. Encouragingly, GL-O15 failed to significantly affect the G4 stability when the G4 had a non-complementary flanking region (Figure S6A). The G4 stabilization effect was regained by altering GL-O15’s guide sequence to match the sequence flanking the G4, GL-O15^ALT^ (Figure S6A-C). Thus, GL-O15 stabilizes only the G4 with a flanking sequence that is complementary to its guide sequence, which is further supported by binding affinity data (Figure S6B-C).

**Figure 4.**
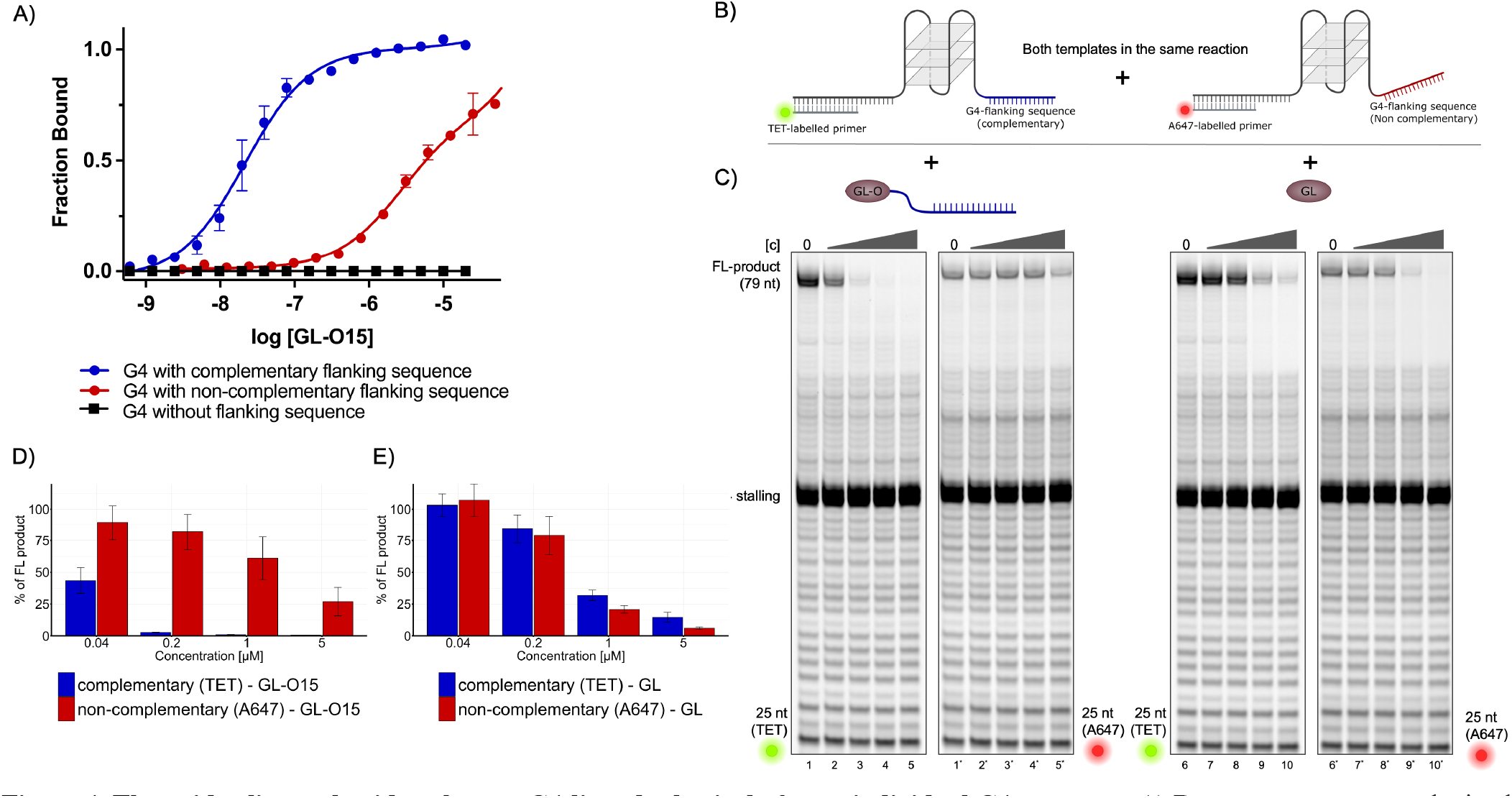
The guide oligonucleotide enhances G4-ligand selectively for an individual G4 structure. **A)** Dose-response curves obtained from MST analysis with GL-O15 using *c-MYC* Pu24T G4 DNA with a flanking sequence that is complementary to GL-O15, a non-complementary flanking sequence, and no flanking sequence. **B)** Schematic representation of the Polymerase-Stop-Assay with two differently labelled templates in the same reaction. Template with a sequence complementary to GL-O15 flanking the G4 was annealed to a 25nt TET labelled primer. Template with an alternative flanking sequence (red) was annealed to a 25 nt A647 labelled primer. Both templates were mixed 1:1 and used for Polymerase-Stop-Assays with **C)** GL-O15 (lanes 2-5 and 2^’^-5^’^) or GL (lanes 7-10 and 7^’^-10^’^) added in increasing concentrations (0.04, 0.2, 1, 5 μM). Left panels of lanes 1-5 and 6-10 represent the scan for the TET label right panels (lanes marked with ^‘^) the scan for A647 of the same lanes. **D)** and **E)** Quantification of the gel shown in C, **D)** for GL-O15 (lanes 2-5 and 2^’^-5^’^) and **E)** for GL (lanes 7-10 and 7^’^-10^’^), percentage of full-length DNA product compared to the control reaction without compound, mean and standard deviation of three individual experiments are shown.

Importantly, GL-O15 is less proficient in stabilizing G4 structures compared to GL when the complementary DNA template sequence is absent (Figure S6A compare lanes 12-15 with 7-10 and 17-20). Combined with the strong binding affinity and G4 stabilization observed from GL-O15 with the correct complementary flanking region (Figure 2D, 3F, S6), this suggests that the guide oligonucleotide sequence of GL-O15 hinders the interaction with the G4 structure in the absence of a complementary sequence on the DNA template, which is of central importance to avoid off-target effects of the approach.

To challenge the specificity, we next performed DNA polymerase stop assays with two competitive G4 containing DNA templates in the same reaction tube, one having a flanking sequence complementary to O15 and another DNA template with an alternative non-complementary flanking sequence (Figure 4B). To distinguish between the DNA products, the non-complementary flanking sequence (which does not form duplex DNA with GL-O15) is A647 labelled, while the DNA substrate with a flanking region complementary to GL-O15 is TET labelled. The control compound (GL) did not discriminate between the two sequences and showed a similar ability to stabilize the G4 structure on both DNA templates (Figure 4C and 4E). In the presence of 1 μM GL, a reduction of about 75% of the fulllength product was detected from both DNA substrates (Figure 4C lanes 9 and 9’). On the contrary, GL-O15 addition displayed an impressive selectivity to the TET labelled DNA template that carries the complementary flanking sequence. A strong reduction (over 90% at 0.2 μM and about 50% at 0.04 μM GL-O15) of full-length product was detected with TET, indicating strong stabilization of the G4 when the complementary flanking region is present (Figure 4C, lanes 2-5, and 4D). The G4 stabilization effect with the non-complementary flanking sequence, however, was really weak as indicated by the modest reduction of the A647 signal (Around 100-1000 times more GL-O15 was needed for the same effect). (Figure 4C, lanes 2’-5’, and 4D). Our experiments have now demonstrated that the GL-O approach result in a synergistic effect that selectively increases the stabilization effect of the target G4 (G4s with a flanking complementary DNA sequence, Figure 3 and S4). Furthermore, the approach actively decreases the unspecific G4 stabilization (non-target G4s with a noncomplementary flanking DNA sequence, Figure 4 and S6). The oligonucleotide conjugation thus specifically guides the GL to the target G4 structure, increases the binding and stabilization of this G4 structure, and prevent off-target interactions.

### The Nucleotide Gap Size Between the G4 Structure and Duplex DNA Formation Exhibits a Degree of Flexibility

The space between the G4 structure and the base pairing location of the oligonucleotide sequence from the GL-O could be critical for its binding and stabilization capacity. To experimentally address this parameter, we synthesized two compounds GL-O16 and GL-O17 in which we increased the length of the guide oligonucleotide fragment compared to GL-O15. The guide oligonucleotide sequence of these molecules was designed to base-pair with the complementary flanking region, 3-nts (GL-O15), 2-nts (GL-O16) or 1-nt (GL-O17) away from the G4 structure (Figure S7A). MST analysis showed that the GL-O16 and GL-O17 derivatives displayed similar binding affinities compared to GL-O15 suggesting that these nucleotide gap alterations tested are not crucial for GL-O binding (Figure S7B-D).

To determine the effect of the nucleotide gap size on G4 stabilization, we performed DNA polymerase stop assays on a *c-MYC* Pu24T containing DNA template that allows hybridization of the oligo fragment of the GL-O15, GL-O16 and GL-O17 compounds. The compounds demonstrated no substantial variation in their capacity to stabilize the G4 structure; when utilizing 0.04 μM of each compound, the reduction of full-length DNA products was consistently around 50% in all instances (Figure S7F and G). This suggests that the GL-O design is not strictly limited to the initially tested 3-nts gap size (between the G4 structure and GL-O hybridization), but there is instead a degree of design flexibility for GL-Os in selecting the guide oligonucleotide sequence based on the G4 flanking region.

### The GL-O Compounds can Differentiate Between G4s with Highly Similar Flanking Regions

We have shown that our GL-O approach does not substantially affect G4s in the absence of any complementary flanking region or with a non-complementary flanking sequence. However, within the human genome there are hundreds of thousands of potential G4 DNA structures. Given this vast number, it is likely that some G4 structures will display sequence homology to the target sequence, potentially influencing the selectivity of our GL-O approach. To investigate this possibility, we altered nucleotides at specific positions of the guide oligonucleotides. All single nucleotide alterations tested did not alter the G4 stabilization ability of GL-O (Figure 5B lanes 2-5 and Figure S8 lanes 7-10, 12-15, 22-25, and 27-30). Similar 2-nucleotide alterations at both ends of the guide oligonucleotide did not considerably affect selectivity (Figure S8 lanes 17-20 and 32-35). In contrast, 2 mismatches in the centre of the guide oligonucleotide greatly influenced the ability of the GL-O (GL-O15 8&11T) to both stabilize and bind the G4 structure (Figure 5). This demonstrates that the GL-O compounds can discriminates against offtarget flanking regions that display great sequence similarity.

**Figure 5.**
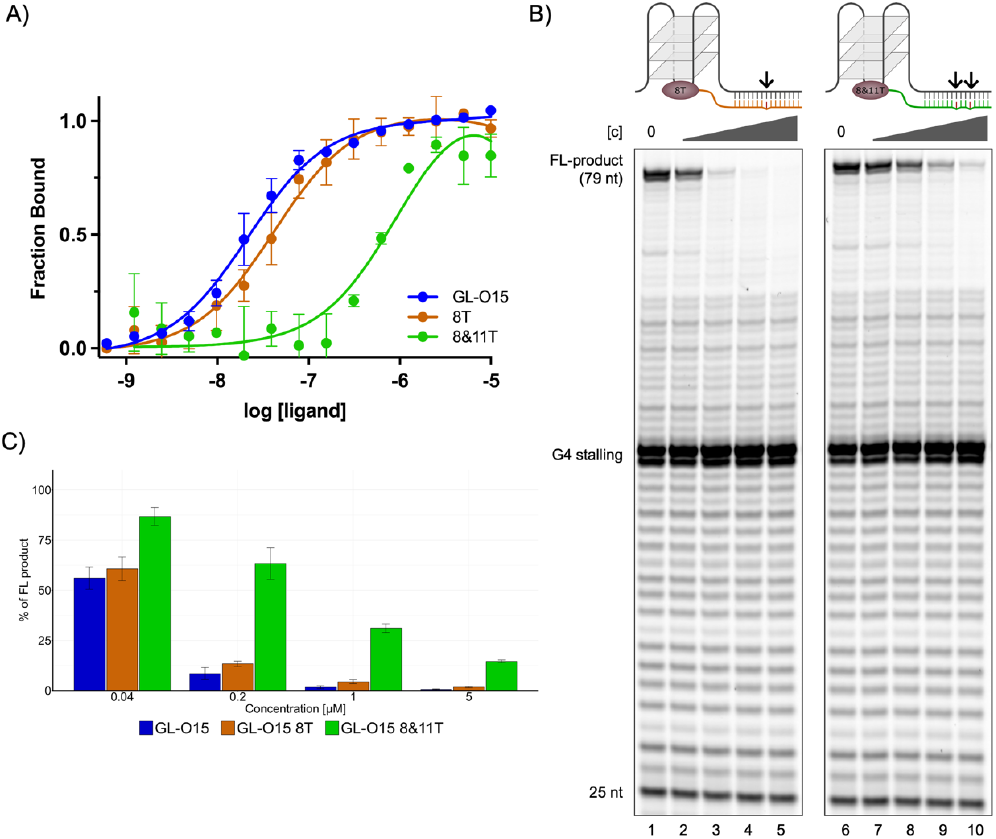
G4 stabilization capability of the GL-O is affected by mismatches in the middle of the oligo. **A)** Dose-response curve obtained from MST analysis after addition of GL-O15, GL-O15 8T and GL-O15 8&11T to *c-MYC* Pu24T G4 DNA with the complementary sequence to GL-O15. Error bars corresponds to SD of two independent measurements. **B)** Polymerase-Stop-Assays in the presence of increasing concentrations (0.04, 0.2, 1, and 5 μM) of GL-O15 8T (lanes 2-5) or with GL-O15 8&11T (lane 7-10). **C)** Quantification of B, percentage of full-length product compared to the control reaction without compound, mean and standard deviation of three individual experiments are shown.

In this work, we developed a strategy based on an unspecific G4-ligand that is conjugated to a DNA oligonucleotide that guides the ligand to only the target G4 structure, leading to selective binding and stabilization of a specific G4 structure. The approach thus combines two recognition devices, one that identifies the correct position in the genome and the other that selectively bind and stabilize G4 DNA. We show that conjugation of the G4-ligand to the guide DNA oligonucleotide does not negatively affect the ability to bind and stabilize G4s. Instead, when conjugated, the G4-ligand and the guide DNA oligonucleotide work in synergy to give higher binding affinities and stabilization than when acting as individual components (Figure 2, Figure 3 and Figure S4). The effect is diminished if only one of them can bind, and off-target effects should therefore largely be avoided. Importantly, we show that the GL-O approach is highly specific to only the target G4 structure and can discriminate between off-target flanking regions with great sequence similarity (Figure 4, S6, and 5). Finally, the distance between the guide sequence and the G4-ligand open for some level of freedom, which can prove to be highly useful in the design of GL-Os to novel targets (Figure S7).

In summary, our presented strategy offers a highly modular approach to specifically target individual G4 structures, which in theory could be used to target any of the hundreds of thousands of potential G4 DNA structures. When considering the diverse regulatory roles of G4s, the presented approach opens for detailed future studies of G4 biology and potential therapeutic interventions. In following studies, we will test our GL-O targeting strategy for individual G4s within a cellular environment.

## MATERIALS AND METHODS

### G4-ligand conjugation of oligonucleotides through click chemistry

A 50 μL of 1 mM oligonucleotide stock was transferred to an Eppendorf to which reagents were added in following order: G4-ligand (compound 6) (5 equiv. 10 mM stock in DMSO/ACN, 3:7), aqueous DIPEA solution (2.5 μl, 0.25 μmol (5 equiv.), 0.043 μl), and CuBr⨯Me2S solution in DMSO (7.5 μl, 0.5 μmol (10 equiv.), 0.1 mg). The reaction mixture was vortexed and subsequently agitated at ambient temperature overnight. The reaction mixture was diluted with 25 μl of an 0.5 mM EDTA solution and 200 μl water before purification by RPHPLC.

### Microscale thermophoresis

G4 DNA labelled with Cy5 at the 5’ end was folded in 10 mM potassium phosphate and 100 mM KCl (pH 7.4) by heating at 95°C for 5 min followed by cooling down to room temperature. All MST experiments were performed in 10 mM potassium phosphate (pH 7.4), 100 mM KCl, 0.05 % Tween 20 and the labelled G4 DNA concentration was held constant at 20 nM. For titration of GL, concentration varied from 0.6 nM to 2.5 μM (thirteen dilution steps). For oligos or ligand conjugated oligos, maximum concentration was fixed at 20 nM and subsequently 1/2 diluted for sixteen steps. The samples were loaded into standard MST grade glass capillaries, and the MST experiment was performed using a Monolith NT.115 (Nano Temper, Germany). For the competition assays 1 equivalent of prefolded Pu24T G4 DNA without flanking sequences or 15-nts were mixed with the Cy5 labelled *c-MYC* Pu24T G4 structure with flanking sequence and MST traces were recorded in presence of different concentrations of oligos or ligand conjugated oligos. MST traces were plotted using OriginPro 8.5 and *K*_*d*_ values were generated by the NanoTemper analysis software. Graphs were further plotted in GraphPad Prism 9.0 for visualization.

### Nuclear magnetic resonance

The G4 DNA stock solution was prepared by folding 100 μM G4 sequences in 10 mM potassium phosphate buffer (pH = 7.4) and 35 mM KCl by heating to 95°C and slowly cooling down to room temperature. 90 μM of effective DNA concentration was obtained by adding 10% D2O to the folded G4 DNA solution. 90 μM of O15/GL-O15 was added to the G4 DNA solution and formation of double standard DNA was monitored. The samples were loaded in 3 mm NMR tubes and ^1^H-NMR spectra were recorded at 298 K on a Bruker 850 MHz Avance III HD spectrometer equipped with a 5 mm TCI cryoprobe. Transmitter frequency offset (O1P) was set at 4.7 PPM and spectral width (SW) was fixed as 22 ppm. Excitation sculpting was used in the 1D ^1^H experiments, and 256 scans were used to record the spectra. For variable temperature NMR, ^1^H-NMR spectra were recorded at 5 °C interval and up to 60 °C. The temperature of the probe was increased manually, and it was allowed to be stable for five minutes before recording the spectra. Processing of the spectra were performed in Topspin 4.1.4 (Bruker Biospin, Germany).

### Circular Dichroism

3 μM of G4 DNA was folded in 10 mM K-phosphate buffer (pH 7.4), with 5 mM KCl by heating for 5 min at 95 °C and then allowed to cool to room temperature. A quartz cuvette with a path length of 1 mm was used for the measurements by JASCO720 spectropolarimeter (Jasco international Co. Ltd.). CD spectra were recorded at 25°C at λ = 210-350 nm with an interval of 0.2 nm and a scan rate of 100 nm/min. Thermal melting curves for G4 DNA were recorded at 263 nm between 20-95°C at a speed of 1 °C/min. Melting temperature (Tm) is defined as the temperature at which 50% of the G4 structures are unfolded.

### Taq Polymerase Stop Assay

The Taq Polymerase Stop assay was adapted from Jamroškovič et al.(19). DNA templates were annealed to fluorescent labelled primers in 100 mM KCl by heating to 95 °C for 5 minutes followed by slow cooling to room temperature (Table S1). The indicated compound concentrations were added to 40 nM annealed template (40 nM of each substrate in the competition assay, Figure 4) in 1x Taq Buffer (10 mM Tris-HCl pH 8.8, 50 mM KCl, Thermo Fisher Scientific), 1.5 mM MgCl_2_, and 0.05 U/μl Taq Polymerase (Thermo Fisher Scientific). Samples were preincubated on ice (10 minutes) and reactions initiated with the addition of dNTPS (100 μM) and transferring the samples to 37 °C. After 15 min at 37 °C reactions were stopped by addition of equal volume of 2x stop solution (0.5% SDS, 25 mM EDTA, XC-Dye in Formamide) and separated on a 12% polyacrylamide Tris-Borate-EDTA (TBE) gel containing 25% formamide and 8 M urea. Fluorescent signal was detected with a Typhoon Scanner (Amersham Biosciences). The intensity of the fulllength band was quantified using Image Quant TL 10.2 software (GE Healthcare Life Sciences) and compared to sample without compound.

## ASSOCIATED CONTENT

## Supporting Information

Supporting tables, figures, synthesis experimental, and NMR Spectra are supplied as supporting information. This material is available free of charge via the Internet at http://pubs.acs.org.

## AUTHOR INFORMATION

## Author Contributions

E.C. S.W. and R.S. conceived the project; A.B., R.N.D., N.B., J.G., A.A., M.A., N.C., M.D., P.P.B, K.C. performed the synthesis/evaluations; E.C., S.W., A.B., R.N.D. N.B. and R.S. analyzed and/or interpreted data; A.B., R.N.D., S.W. and E.C. wrote the manuscript. All authors have given approval to the final version of the manuscript. ^#^ Equally contributing first authors, ^*^ Equally contributing corresponding authors.

### Funding Sources

Work in the Chorell lab was supported by the Kempe Foundations (JCK-3159 and SMK-1632) and the Swedish Research Council (VR-NT 2017-05235 and VR-NT 2021-04805). Work in the Wanrooij lab was supported by the Knut and Alice Wallenberg Foundation, Kempe Foundations (SMK21-0059) and the Swedish Research Council (VR-MH 2018-0278).

---

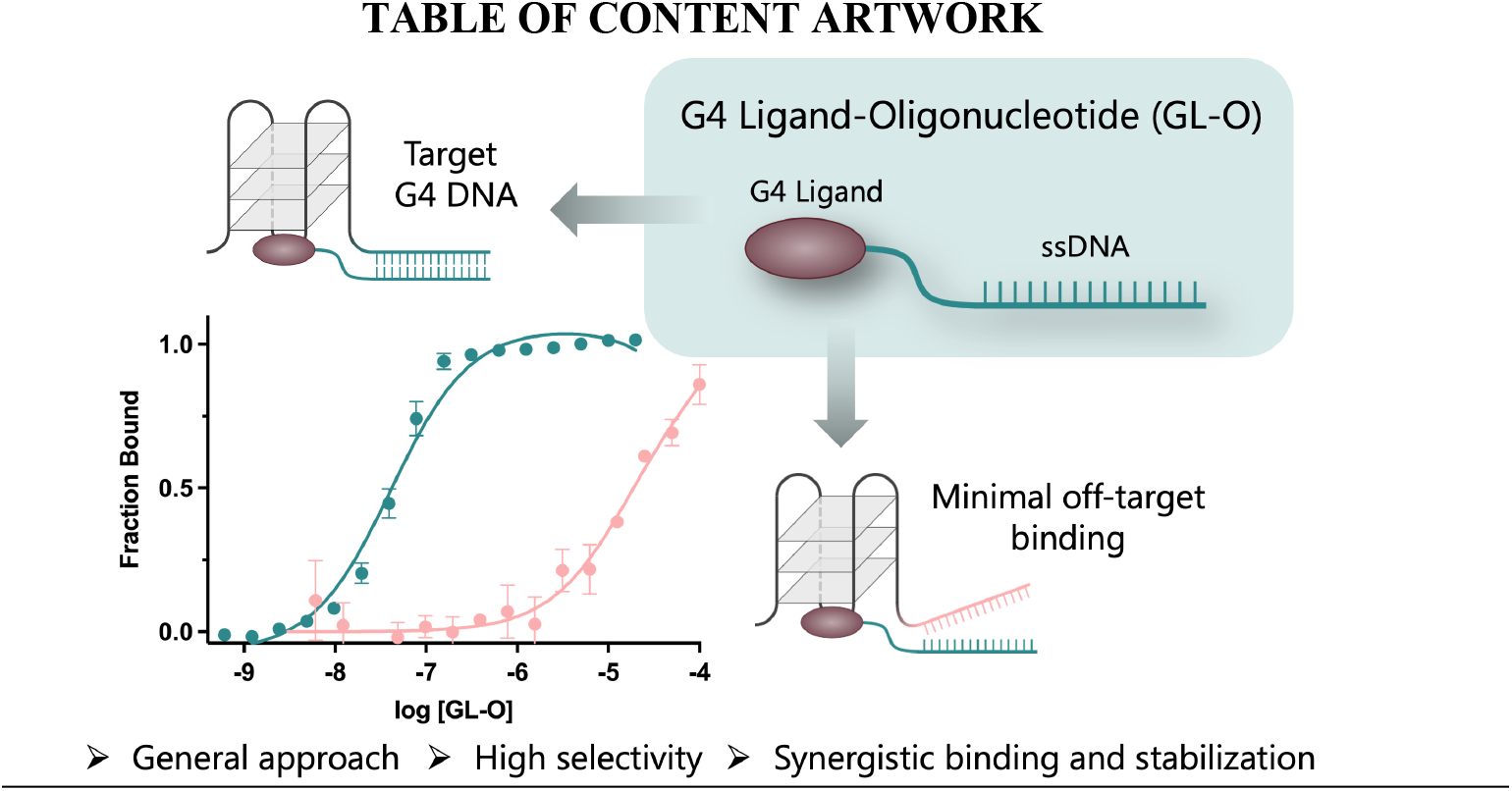

